# Structural context modulates the conformational ensemble of the intrinsically disordered amino terminus of α-synuclein

**DOI:** 10.1101/2024.10.31.621304

**Authors:** Rania Dumarieh, Dominique Lagasca, Sakshi Krishna, Jaka Kragelj, Yiling Xiao, Kendra K. Frederick

## Abstract

Regions of intrinsic disorder play crucial roles in biological systems, yet they often elude characterization by conventional biophysical techniques. To capture conformational distributions across different timescales, we employed a freezing approach coupled with solid-state NMR analysis. Using segmentally isotopically labeled α-synuclein (α-syn), we investigated the conformational preferences of the six alanines, three glycines, and a single site (L8) in the disordered amino terminus under three distinct conditions: in 8 M urea, as a frozen monomer in buffer, and within the disordered regions flanking the amyloid core. The experimental spectra varied significantly among these conditions and deviated from those of a statistical coil. In 8 M urea, monomeric α-syn exhibited the most restricted conformational sampling, rarely accessing chemical shifts characteristic of α-helices or β-strands. In buffer, monomeric α-syn showed broader conformational sampling, favoring α-helical conformations and, to a lesser extent, random coil states. Notably, amino acids in the disordered regions flanking the amyloid core demonstrated the most extensive conformational sampling, with broad peaks encompassing the entire range of possible chemical shifts and a marked preference for highly extended β-strand conformations. Collectively, this work demonstrates that intrinsically disordered regions exhibit distinct conformational preferences, which are influenced not only by the chemical environment but also by the conformations of adjacent protein sequences. The differences in the conformational ensembles of the disordered amino terminus may explain why the monomer and the amyloid form of α-syn interact with different biomolecules inside cells.

## INTRODUCTION

The structural characterization of intrinsically disordered proteins demands innovative approaches that can capture their dynamic, context-dependent conformational preferences.(1, 2). The protein α-synuclein (α-syn) exemplifies the biological significance of intrinsic disorder, as its conformational plasticity is central to both its physiological function and its role in neurodegenerative disease (3). In its monomeric form, α-syn is completely intrinsically disordered (4), but it can adopt diverse conformational states depending on its environment. These include α-helical rich structures in the presence of lipids and various β-sheet-rich amyloid fibril forms that differ in the number and arrangement of β-sheets (5, 6). The aggregation of α-syn into amyloid fibrils is a hallmark of several neurodegenerative disorders. In these fibrillar structures, while the central ∼70 amino acids form an ordered amyloid core through serpentine arrangements of β-strands and connecting turns, both the amino terminal (∼30 residues) and carboxy terminal (∼40 residues) regions remain disordered, creating a "fuzzy coat" around the fibril core. Understanding the conformational ensemble of these persistently disordered regions is crucial for elucidating both the physiological function of α-syn and its role in disease pathogenesis.

Intrinsically disordered regions (IDRs) display distinct characteristics based on their composition. Sequence-encoded features, such as net charge per residue, modulate their conformational ensembles (7, 8) as do various co-solutes (9). However, the influence of adjacent amino acid conformations on these ensembles remains less understood. While NMR studies have confirmed that amino terminus of α-syn is disordered in both monomeric and fibrillar forms, monomers and fibrils interact with different biomolecules inside cells. Moreover, these flexible regions mediate primary and secondary nucleation (10–12) and facilitate interactions between amyloid fibers and cellular constituents, particularly chaperone proteins (13–17). While the flanking regions significantly influence aggregation, toxicity, and cellular (dys)function (18, 19), their structural details remain elusive. Understanding the conformational preferences of these disordered regions could reveal new therapeutic targets for aggregation-related diseases (20, 21).

Solution-state NMR chemical shifts report on conformationally weighted averages for sites interconverting on nanosecond timescales, making NMR a valuable tool for validating molecular dynamics (MD) ensembles (22) of intrinsically disordered regions. Indeed, solution state NMR experiments indicate that monomeric α-syn is intrinsically disordered and preferentially samples α-helical dihedral angles (4). In contrast, dipolar-based solid-state NMR reports on sites that are immobile on microsecond or longer timescales; mobile sites are not observed. In solid state NMR experiments of an amyloid form of α-syn at room temperature, residues 38-96 were sequestered in the rigid amyloid core (23, 24) while directly flanking regions (residues 30-38 and 104-110) experienced intermediate time scale motions and amino and carboxy termini that flanked the amyloid core were not seen due to their intrinsically disordered nature (24).

One approach to capture conformational distributions across different timescales involves freezing samples for solid-state NMR analysis (25–29). While cross-peak line shapes in frozen samples have long served as quantitative constraints on backbone conformational distributions (25–29) recent advances in DNP MAS NMR, including higher magnetic fields and improved cryogenic probes, have enhanced spectral resolution. For highly ordered systems, line widths at ambient and cryogenic temperatures remain similar (30, 31). This indicates that the broader line widths at cryogenic temperatures reflect frozen local conformational fluctuations typically undetected at room temperature (32–36). This effect is exaggerated for loops and intrinsically disordered regions because the peak reports on all the sampled conformations with peak intensities related to the relative populations of each conformation in the ensemble (37).

We recently developed a framework for obtaining experimental conformational restraints in frozen samples through peak shape analysis (37). This approach compares experimental spectra to those predicted for unrestrained amino acids in a polypeptide chain, highlighting conformational preferences relative to statistical coil conformations. Here, we combine this method with segmental and specific isotopic labeling strategies using intein-mediated protein ligation (38–41). Inteins, self-excising amino acid sequences that ligate flanking sequences via native peptide bonds (42), specifically the Cfa intein (43), enable site-specific isotopic labeling. Here, we desulfurize of the inserted catalysitc cysteine to alanine (44) to create a mutationless segmentally isotopically labeled protein(41). We apply this technique to examine α-syn’s amino terminus in various states: urea-denatured, as an intrinsically disordered monomer in buffer, and within the disordered "fuzzy coat" of an amyloid fibril (23). By comparing these experimental spectra with statistical coil predictions, we investigate how environment and proximal structural constraints influence these intrinsically disordered regions.

## RESULTS

There are six alanine residues – A11, A17, A18, A19, A27, and A29 – in the amino terminal sequence of α-syn (**Figure 1**). The predicted shape for the Cα-Cβ cross peak from an ensemble of statistical coil conformations of alanine spanned a wide range of chemical shifts (**Figure 1A**) and was fit to four Gaussian peaks with an average FWHM of ∼2.4 ppm ± 0.6 ppm (**Table S1**). One peak was centered at 55.3 ppm and 18.0 ppm, for the Cα and the Cβ, respectively, near the average chemical shift for α-helical dihedral angles and accounted for 16% of the population. Another was centered at 53.25 ppm and 18.6 ppm near the average chemical shift for random coil conformations and accounted for 60% of the population. A third was centered at 51.3 ppm and 21.0 ppm near the average chemical shift for β-strands and accounted for 15% of the population. Finally, there was a peak centered at 53.3 ppm and 16.0 ppm, a region that maps to conformations with positive phi angles, which accounted for 8% of the population (**Figure 1A, Table S1**). The alanine cross-peak is well resolved from the cross peaks for other amino acids. To experimentally characterize the conformational ensemble for the six alanines in the amino terminal region of α-syn, we compared the chemical shifts and the integrated peak intensities for the Cα-Cβ cross-peak of alanine of frozen samples of segmentally labeled α-syn in three forms: as a chemically denatured monomer, as a monomer in buffer, and in the disordered region that flanks the amyloid fibril form. These were compared to each other and to the predicted peak shape for a statistical coil.

**Figure 1:**
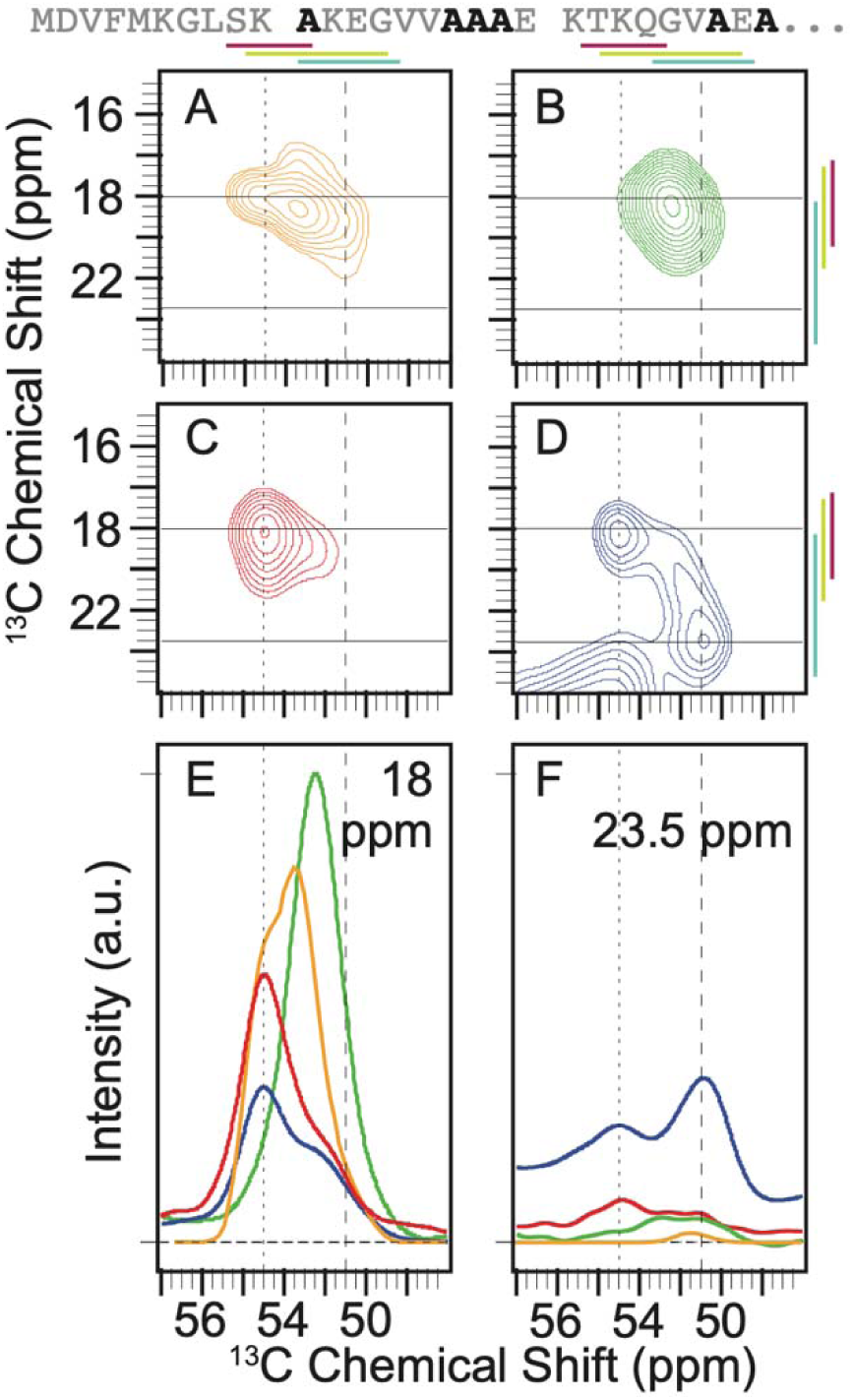
Context alters the conformational preferences of alanine residues in the amino terminal intrinsically disordered region of α-syn. The primary sequence of the isotopically labeled segment of α-syn is shown with alanine residues highlighted in black. A) The predicted peak shape for the Cα-Cβ cross peak of alanine from a statistical coil structural ensemble assuming a homogenous line width of 1.5 ppm. ^13^C-^13^C DARR spectra of frozen segmentally isotopically labeled α-syn in B) 8M urea C) in the monomeric form in buffer and D) in the amyloid fibril form in buffer. Colored bars annotate the average chemical shift ± two standard deviations for α-helices (magenta), random coils (light green) and β-strands (light blue) (45). Spectra were normalized to have the same integrated intensity. Horizontal lines at 18 ppm and 23.5 ppm mark the location of 1D slices in E and F. E) Overlay of the normalized one-dimensional slices from the 2D DARR spectra at 18 ppm. F) Overlay of normalized one-dimensional slices from the 2D DARR spectra at 23.5 ppm. Samples contained 15% d_8_-glycerol and 5 mM AMUPol. Spectra were collected at 600 MHz with 12 kHz MAS at 104 K.

In 8 M urea, the Cα-Cβ cross peak for alanine sampled a smaller range of chemical shifts than the statistical coil prediction. The cross peak fit to a single Gaussian centered at 52.4 ppm and 18.5 ppm with a FWHM of ∼4 ppm in both dimensions (**Figure 1B**). In buffer, where long range interactions are possible, frozen monomeric α-syn sampled more conformational space than in 8 M urea and had a marked preference for α-helical chemical shifts (**Figure 1C**). The peak for the frozen monomer in buffer was fit to two Gaussians. One Gaussian was centered at 55.0 ppm and 18.4 ppm, consistent with α-helical dihedral angles, and accounted for two thirds (67% ± 1%) of the peak intensity. The second Gaussian was centered at 52.2 ppm and 19.4 ppm, consistent with extended coil conformations and accounted for the other third of the intensity (33% ± 1%). When α-syn is assembled into amyloid fibrils, the amino terminal 38 amino acids are mobile on the microsecond or faster timescales (23) and are therefore assumed to be intrinsically disordered. The peak for the alanine residues in the flexible flanking region of the amyloid fibril was described by three Gaussians (**Figure 1D**). The intensity was split evenly between all three broad peaks (FWHM 3.2 ppm ± 0.8 ppm, **Table S1**). The peak that corresponded to α-helical dihedrals was centered at 55.1 ppm and 18.5 ppm and accounted for 28% ± 1% of the population. Like the frozen monomeric α-syn in buffer, the peak consistent with extended coil conformation was centered at 52.3 ppm and 18.9 ppm and accounted for about a third of the intensity (34% ± 6%). The Gaussian that was centered at 50.8 ppm and 23.3 ppm, which is consistent with β-strands, accounted for 38% ± 7% of the population.

Interestingly, the average chemical shift for the intrinsically disordered flanking region was shifted towards the extreme end of the possible values for extended conformations (**Figure 1**, light blue lines) (45). Moreover, while dihedrals consistent with β-strand conformations represented about 15% of the population of the statistical ensemble, such conformations were sampled by the intrinsically disordered flanking region of α-syn more than twice as frequently. These more extreme chemical shift values were rarely observed in any of the other examined experimental or theoretical contexts (**Figure 1F**). Overall, we found being adjacent to the amyloid core profoundly altered the conformational ensemble of the flanking intrinsically disordered regions of the amyloid fibril form.

There are three glycine residues – G7, G14, and G25 – spaced across the amino terminal sequence of α-syn (**Figure 2**). The predicted peak shape from an ensemble of statistical coil conformations was well described by a single broad Gaussian peak with a FWHM of ∼3 ppm centered at 44.7 and 174.2 ppm for the Cα and CO, respectively, which is consistent with random coil conformations (**Figure 2A, Table S1**). To experimentally characterize the conformational ensemble for the glycines in the amino terminal region of α-syn, we compared the chemical shifts and the integrated peak intensities for the Cα-CO cross-peak of glycine, which is well resolved from the cross peaks for other amino acids, for frozen samples of segmentally labeled α-syn of a chemically denatured monomer, as a monomer in buffer and for α-syn in the amyloid fibril form to each other and to the predicted peak shape for a statistical coil. In 8 M urea, the glycine peak is well described by a single broad Gaussian peak centered at 44.6 and 174.3 ppm with a FWHM of 4 ppm (**Figure 2B**). The experimental spectra for the chemically denatured sample, where urea eliminated long range interactions, and the spectra for a statistical coil ensemble were centered at the same position, but the experimental spectrum was broader. The broader spectrum indicates that the most highly populated conformations are similar in these two ensembles, but the glycines in this region of α-syn in 8 M urea sample a higher proportion of conformational space than described by the statistical coil approximation.

**Figure 2:**
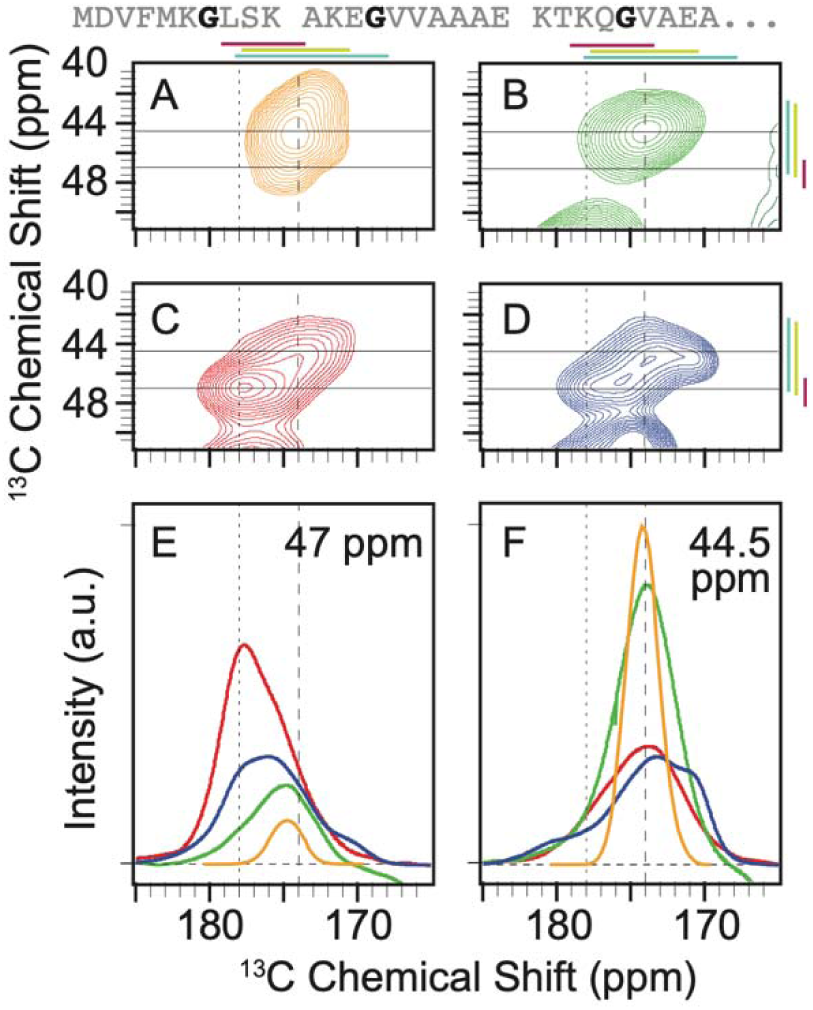
Context alters the conformational preferences of glycine residues in the amino terminal intrinsically disordered region of α-syn. The primary sequence of the isotopically labeled segment of α-syn is shown with glycine residues highlighted in black. A) The predicted peak shape for the Cα-CO cross peak of glycine from a statistical coil structural ensemble assuming a homogenous line width of 1.5 ppm. ^13^C-^13^C DARR spectra of frozen segmentally isotopically labeled α-syn in B) 8 M urea C) in the monomeric form in buffer and D) in the amyloid fibril form in buffer. Colored bars annotate the average chemical shift ± two standard deviations for α-helices (magenta), random coils (light green) and β-strands (light blue). Spectra were normalized to have the same integrated intensity. Horizontal lines at 47 ppm and 44.5 ppm mark the location of 1D slices in E (47 ppm) and F (44.5 ppm). E) Overlay of the normalized one-dimensional slices from the 2D DARR spectra at 47 ppm. F) Overlay of normalized one-dimensional slices from the 2D DARR spectra at 44.5 ppm. Samples contained 15% d_8_-glycerol and 5 mM AMUPol. Spectra were collected at 600 MHz with 12.5 kHz MAS at 104 K.

To determine the ensemble of conformations sampled by glycine in the amino terminal region of α-syn in a context where long range transient interactions can alter the conformational ensemble, we froze monomeric α-syn in buffer. We found that for the frozen monomer in buffer, the intensity of the glycine cross peak spans a wider range of chemical shifts than in 8 M urea (**Figure 2C**). Like the alanine residues for the monomer in buffer, the glycines preferentially adopted the dihedral angles found in α-helices. The peak for the frozen monomer in buffer was fit to three Gaussians, each with FMWH of ∼4 ppm (**Table S1**). The Gaussian centered at a position consistent with α-helical dihedral angles (47.0 ppm and 177.6 ppm) accounted for half (55% ± 3%) of the intensity. Most of the remaining the population (40% ± 4%) was described by a Gaussian centered at 45.6 and 174.7 ppm, consistent with random coil conformations. A small population (5% ± 2%) was described by a Gaussian centered at 43.8 ppm and 171.9 ppm, consistent with extended β-strands. Therefore, the intrinsically disordered monomer preferentially sampled dihedral angles consistent with α-helices.

Interestingly, when α-syn was assembled into amyloid fibrils, while the conformations adopted by the glycines sample a similar range of conformational space to that of the monomer in buffer, the relative populations of the conformations were different (**Figure 2D**). The peak that corresponded to α-helical dihedrals was centered at 46.9 ppm and 177.5 ppm but was half as populated as in the monomer (26% ± 3%). The peak consistent with extended coil conformation at 45.0 ppm and 173.7 ppm accounted for more than half of the intensity (55% ± 1%). The population center that corresponds to extended β-strands was shifted to more extreme values at 44.9 and 170.6 ppm and was three times as populated (18% ± 2%) as the same region of α-syn in the monomeric form. The spectrum of the flanking disordered region of the amyloid fibril indicated that glycines in this disordered region adopt highly extended conformations like those found in β-strands twice as often as when α-syn is in the monomeric form.

There is only a single leucine residue – L8 – in the disordered amino terminal region of α-syn (**Figure 3**). The predicted shape for the Cα-Cβ cross peak from an ensemble of statistical coil conformations for leucine spanned a wide range of chemical shifts (**Figure 3A**). It was fit to four Gaussian peaks with an average FWHM of ∼2.2 ppm ± 0.6 ppm. One Gaussian was centered at 56.9 ppm and 41.2 ppm for the Cα and Cβ, respectively, near the average chemical shift for α-helical dihedral angles and accounted for 8% of the population. Another Gaussian was centered at 54.5 ppm and 41.6 ppm near the average chemical shift for random coil conformations and accounted for 64% of the population. A third Gaussian was centered at 52.4 ppm and 42.7 ppm near the average chemical shift for β-strands and which accounted for 21% of the population. The final Gaussian was centered at 53.7 ppm and 39.2 ppm, a region that maps to conformations with positive phi angles and accounted for 7% of the populations (**Figure 3A, Table S1**).

**Figure 3:**
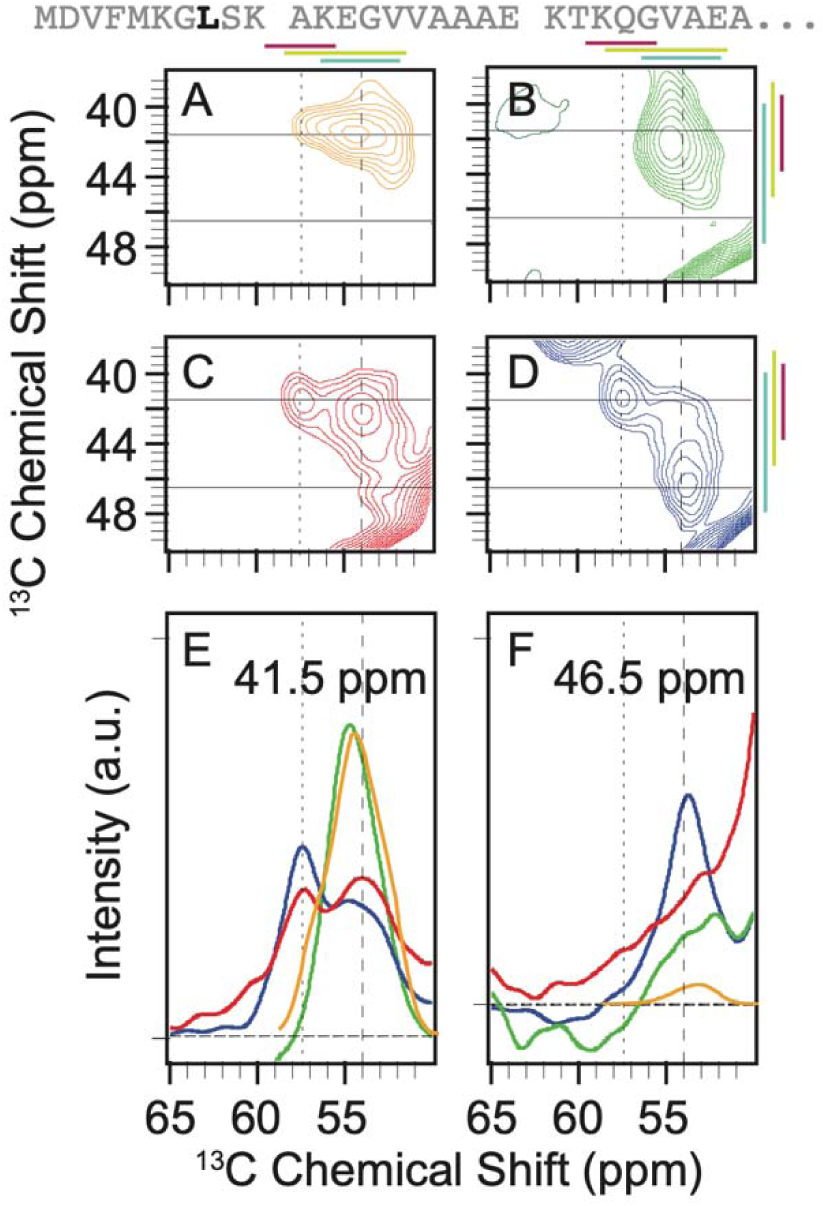
Context alters the conformational preferences of leucine 8 of α-syn. The primary sequence of the isotopically labeled segment of α-syn is shown with leucine residues highlighted in black. A) The predicted peak shape for the Cα-Cβ cross peak of leucine from a statistical coil structural ensemble assuming a homogenous line width of 1.5 ppm. ^13^C-^13^C DARR spectra of frozen segmentally isotopically labeled α-syn in B) 8M urea C) in the monomeric form in buffer and D) in the amyloid fibril form in buffer. Colored bars annotate the average chemical shift ± two standard deviations for α-helices (magenta), random coils (light green) and β-strands (light blue). Spectra were normalized to have the same integrated intensity. Horizontal lines mark the location of 1D slices in E and F. E) Overlay of the normalized one-dimensional slices from the 2D DARR spectra at 41.5 ppm. F) Overlay of normalized one dimensional slices from the 2D DARR spectra at 46.5 ppm. Samples contained 15% d_8_-glycerol and 5 mM AMUPol. Spectra were collected at 600 MHz with 12 kHz MAS at 104 K.

In 8 M urea, the leucine peak sampled less conformational space than the statistical coil prediction and was described by a single broad Gaussian peak centered at 54.7 ppm and 41.0 ppm with a FWHM of 3 ppm for the Cα and 4 ppm for the Cβ (**Figure 3B**). In buffer, L8 sampled more conformational space than in 8 M urea and preferentially adopted α-helical dihedral angles. The peak for the frozen monomer in buffer was fit to two Gaussians (**Figure 3C**). The Gaussian centered at a position consistent with α-helical dihedral angles (57.4 ppm and 41.3 ppm), accounted for almost half (40% ± 1%) of the intensity. The rest of the population (60% ± 1%) was described by a Gaussian centered at 54.0 ppm and 42.6 ppm, consistent with random coil conformations. Therefore, like the alanine and glycine residues, L8 in the intrinsically disordered monomer preferentially adopted dihedral angles consistent with α-helices. When α-syn was assembled into amyloid fibrils, the leucine peak covered a wider range of conformational space than in the monomeric form (**Figure 3D**). The peak that corresponds to α-helical dihedrals was centered at 57.5 ppm and 41.3 ppm and was less populated than in the monomer in buffer (30% ± 1%). The peak consistent with extended coil conformation was centered at 54.1 and 42.6 ppm was half as populated as in the monomeric state (31% ± 1%). The region that corresponds to extended β-strands was centered at ppm 53.7 ppm and 46.5 ppm and became the most populated (41% ± 1%) of the three general peak regions we considered. The statistical coil approximation did not capture the distribution of conformations of L8 in any of the examined experimental contexts. L8 of α-syn in 8 M urea samples the smallest range of backbone dihedral conformations, followed by frozen monomer of α-syn in buffer, which samples a broader range with a preference for a-helical conformations. Finally, as observed for the alanine and glycine residues, L8 in the intrinsically disordered region that flanks the amyloid core sampled the largest range of conformations, with a marked preference for extended β-strands conformations. Unlike the alanines and glycines, there is only one leucine in this region. While the broader distributions in the fuzzy coat could derive from different residues preferentially sampling different conformations within the same strand for alanine and glycine, that is not the case for leucine. Therefore, the individual strands in the fuzzy coat adopt a variety of distinct conformations.

Serine and threonine residues are each represented only one time in the amino terminal region of α-syn and the serine Cα-Cβ and threonine Cα-Cβ cross peaks that have chemical shifts that are well-resolved from those of other amino acids. However, the chemical shifts for these residues merge with the diagonal feature of the spectrum. Thus, the conformational preferences can only be partially assessed for S9 and T22. Nonetheless, a partial analysis indicated that the conformational preferences at these sites resemble those observed for the alanines, glycines and L8 across the studied conditions.

The predicted shape for serine Cα-Cβ cross peak from an ensemble of statistical coil conformations for threonine spanned a wide range of chemical shifts (**Figure S1**), most conformations result in cross-peaks near the diagonal feature of the spectrum. The chemical shifts for extended β-strand conformations are the furthest from the diagonal (**Figure S1,** bars). In the experimental spectra, the S9 peak was merged with the diagonal feature. The only spectrum that had a distinct feature was that of S9 in the disordered region that flanked the amyloid core. This spectrum had a peak consistent with a population that sampled extended β-strand conformations (**Figure S1D, F**).

The predicted shape for threonine Cα-Cβ cross peak from an ensemble of statistical coil conformations for threonine spanned a wide range of chemical shifts (**Figure S2**), many of which result in cross-peaks near the diagonal feature of the spectrum (**Figure S2F**). The predicted shape for threonine Cα-CO cross peak from an ensemble of statistical coil conformations for threonine likewise spanned a wide range of chemical shifts, however those for α-helical conformations were not degenerate with chemical shifts for other amino acid types present in the sample (**Figure S2K**). In contrast, while the Cβ-Cg cross-peak is degenerate with other amino acids for α-helical conformations, it is not degenerate with other amino acids for β strand and coil conformations (**Figure S2A**). With these caveats, in 8 M urea, T22 did not populate either α-helical (**Figure S2M**) or β-strand (**Figure S2C**) conformations. For monomers frozen in buffer, T22 populated α-helical conformations (**Figure S2B, L**) but did not populate extended beta conformations (**Figure S2G, B**). Finally, T22 in the disordered region that flanked the amyloid core populated both α-helical (**Figure S2D, N**) and extended β-strand populations (**Figure S2D, E, I**). Moreover, as seen for other amino acid types, T22 preferentially sampled α-helical conformations in the frozen monomeric form in buffer and β-strands in the disordered regions that flank the amyloid core (**Figure S2E, J, O**). Thus, qualitatively, the changes in the conformational ensemble for T22 across the examined conditions resembles the changes observed for alanines, glycines, and L8.

## DISCUSSION

Intrinsically disordered proteins challenge traditional structural biology paradigms by presenting dynamic, context-dependent conformational landscapes that resist static structural characterization. To capture conformational distributions across different timescales, we employed a freezing approach coupled with solid-state NMR analysis. Using segmentally isotopically labeled α-syn, we investigated the conformational preferences of the six alanines, three glycines, and a single site, L8, in the disordered amino terminus under three distinct conditions: in 8 M urea, as frozen monomer in buffer, and in the disordered regions flanking the amyloid core. The experimental spectra differed from each other as well as from that of a statistical coil. In 8 M urea, monomeric α-syn exhibited the most restricted conformational sampling, rarely accessing chemical shifts characteristic of α-helices or β-strands. Monomeric α-syn in buffer had broader conformational sampling, with a preference for α-helical conformations and, to a lesser extent, random coil conformations. Notably, amino acids in the disordered regions flanking the amyloid core demonstrated the most extensive conformational sampling, with broad peaks that covered the entire range of possible chemical shifts with a marked preference for highly extended β-strand conformations. Collectively, this work demonstrates that intrinsically disordered regions have conformational preferences and that these conformational preferences are altered by not only the chemical environment but also conformation of the adjacent protein sequence.

Several experimental constraints warrant consideration when interpreting our results. Although all samples were frozen uniformly (cooling from room temperature to 100 K in approximately 10 seconds), this rate remains substantially slower than most molecular motions, which can potentially introduce conformational bias during the freezing process. Additionally, our experimental requirement for vitreous ice necessitated the inclusion of 15% glycerol as a cryoprotectant to prevent ice crystal formation. The impact of glycerol on the conformational ensemble remains undetermined. Moreover, from an information theory perspective, our experiments provide primarily local conformational information through site-specific chemical shifts. While these measurements effectively report on the distribution of backbone dihedral angles at individual positions, they cannot capture the mutual information between spatially separated regions of the protein. Such correlated motions, which can arise from both through-space interactions and through-backbone propagation of conformational preferences, are particularly relevant in intrinsically disordered proteins where transient long-range contacts can significantly influence the overall conformational ensemble. Moreover, the frozen-state nature of our measurements precludes direct information about the timescales and coupling of these correlated motions at room temperature, where dynamic processes may link seemingly independent local conformational changes. Despite these limitations, the consistent sample preparation and analysis conditions across all experimental states enable meaningful comparative analysis. While the absolute values we observe may not precisely reflect room temperature ensembles, the observed differences between conformational distributions under various conditions represent genuine variations in conformational preferences.

Our comparison of intrinsically disordered monomeric α-syn in 8 M urea with statistical coil predictions reveals an apparent paradox in conformational sampling. The strong chaotropic environment stabilizes a more extended conformation (9, 46) while simultaneously restricting the conformational space that is meaningfully sampled. Counterintuitively, what is commonly considered the most "disordered" state exhibits the least conformational diversity from a statistical mechanics perspective. This observation can be reconciled by considering two competing effects of chaotropes on intrinsically disordered proteins. On one hand, chaotropes disrupt hydrogen bonds and eliminate hydrophobic collapse, (46–48) preventing long-range interactions that would otherwise constrain the conformational ensemble. This effect should drive the 8 M urea state toward statistical coil behavior. However, the increased effective size of the protein-solvent system in concentrated urea solutions sterically restricts access to certain backbone dihedral angles, particularly those that are frequently sampled in aqueous conditions (49, 50). The net result is a conformational ensemble that, despite the absence of stabilizing intramolecular interactions, samples fewer states than predicted by statistical coil models. These findings highlight how solvent properties—both size and chemistry—can significantly modulate the conformational landscape of disordered proteins.

Our analysis of α-syn in frozen buffer reveals significant deviations from statistical coil predictions, despite the expected solvent properties of water. These differences likely arise because the statistical coil ensemble do not account for sequence context or longer-range interactions (37). Notably, we observed that the population of α-helical chemical shifts for alanine, glycine, L8, and T22 all exceed statistical coil predictions by more than twofold. This enhanced α-helical propensity aligns qualitatively with room-temperature solution-state NMR experiments, which have established that monomeric α-synuclein displays inherent α-helical preferences in solution (4). However, the α-helical bias we observe for these sites is more pronounced than what might be predicted from the time-averaged chemical shifts in solution-state NMR experiments (51). While this heightened helical propensity could be an artifact of our experimental conditions, it may alternatively reveal conformational states that are sampled on timescales beyond the detection limits of solution-state NMR (10, 52).

The disordered region that flanks the amyloid core has in a unique chemical environment, where individual polypeptide chains are constrained at one end in a regular array with 4.7 Å spacing as they emerge from the ordered amyloid core. This arrangement creates a locally crowded environment with polyvalent characteristics, significantly influencing conformational sampling. Our analysis reveals that this region samples α-helical conformations approximately 1.5-fold more frequently than statistical coil predictions, while extended β-strand conformations are enriched two- to three-fold. These observations complement room temperature solid-state NMR studies, where residues 1-38 are absent, indicating that the flanking region is dynamic on microsecond or faster timescales (23). Indeed, while residues 1-29 are not detected, NMR experiments sensitive to motions on the 100’s of millisecond timescale detect residues 30-38, which have chemical shifts consistent with β-strands (24). Interestingly, however, ordered states and motifs have been reported in the regions that flank the amyloid core of α-syn (10, 20, 53). Therefore, particularly in the context of ordered states, the substantial conformational heterogeneity we observe warrants careful interpretation. For alanine and glycine, the broad conformational distribution could simply reflect different sites adopting distinct stable conformations. However, site-specific measurements provide compelling evidence for genuine conformational heterogeneity. T22, positioned 16 amino acids from the amyloid core and 8 amino acids from the last known β-sheet-prone region, samples both α-helical and β-strand conformations. More striking is L8, located 30 amino acids from the core, which shows a marked preference for extended β-strand conformations while still sampling a broad conformational space. The conformational preferences of these individual sites mirror the distributed sampling observed for the alanine and glycine ensembles, confirming that neither L8 nor T22 adopts a single stable conformation.

## Acknowledgements

D.L. was supported by NIH MB T32 GM008297. S.K. was supported by UT Dallas and the Cecil and Ida Green Foundation via The Green Fellow’s Program. This work was supported by grants from the National Institute of Health [NS111236 and NS134921 and R21NS136951] to K.K.F.

## Data and materials availability

Code to generate statistical coil ensembles is available at https://github.com/Sakshi-Krishna/FrederickLab/tree/main/Predictions

## MATERIALS AND METHODS

### Split-Intein Construct Preparation

The genes for the split intein Cfa_GEP_ and α-syn were codon-optimized for bacterial expression by GenScript. Construct 1 contained the first 29 amino acids of α-syn followed by the first 101 amino acids of the Cfa_GEP_ sequence and was cloned into the pET-28a(+) vector (Invitrogen). Construct 2 contained the last 36 amino acids of the Cfa_GEP_ intein and residues 30 to 140 of α-syn (with alanine 30 mutated to cysteine) and was cloned into the NcoI and XhoI sites of the pET-28a(+) vector using the N-terminal hexahistidine tag encoded by the vector.

#### Construct 1 Sequence

α-syn(1–29)-<u>Cfa_N_</u>-linker-<u>Thiredoxin</u>-linker-<u>His_6_</u>MDVFMKGLSK AKEGVVAAAE KTKQGVAEA<u>C LSYDTEILTV EYGFLPIGKI VEERIECTVY TVDKNGFVYT QPIAQWHNRG EQEVFEYCLE DGSIIRATKD HKFMTTDGQM LPIDEIFERG LDLKQVDGLP</u> SSGPS<u>MSDKI IHLTDDSFDT DVLKADGAIL VDFWAEWCGP CKMIAPILDE IADEYQGKLT VAKLNIDQNP GTAPKYGIRG IPTLLLFKNG EVAATKVGAL SKGQLKEFLD ANLA</u>GAASG<u>H HHHHH</u>

#### Construct 2 Sequence

His_6_-linker-caspase 3 cleavage site-<u>Cfa_C_</u>-α-syn(C30-140) MGSDKIHHHH HH<u>SSGGSGAA SG</u>GDEVD<u>MVK IISRKSLGTQ NVYDIGVGEP HNFLLKNGLV ASN</u>**C**GKTKEG VLYVGSKTKE GVVHGVATVA EKTKEQVTNV GGAVVTGVTA VAQKTVEGAG SIAAATGFVK KDQLGKNEEG APQEGILEDM PVDPDNEAYE MPSEEGYQDY EPEA

### Protein Expression and Purification

To express the isotopically-labeled recombinant α-syn(1–29)-Cfa_N_ construct, an overnight pre-culture of BL21(DE3) cells carrying the plasmid encoding syn(1–29)-Cfa_N_ was used to inoculate 4 liters of minimal media containing 50 mg/L kanamycin (composition of minimal media detailed below). Cells were grown at 37 °C with shaking until their *A*_600_ reached 0.6–0.8; the protein expression was then induced by adding isopropyl 1-thio-β-D-galactopyranoside (IPTG) to a final concentration of 1 mM. Cells were harvested 4 hours after induction by centrifugation at 4000 × *g* for 15 min at 4 °C. For ^13^C, ^15^N-segmentally labeled synuclein, minimal media contained the following: M9 salts (48 mM Na_2_HPO_4_, 22 mM KH_2_PO_4_, 9 mM NaCl, adjusted to pH 7.4 by the addition of NaOH), 4 g/L ^13^C_6_-glucose, 1 g/L ^15^NH_4_Cl, 10 mg/L FeSO_4_, 2 mM MgSO_4_, 100 µM CaCl_2_, 10 mg/L thiamine). For ^13^C, ^15^N-VGLAST-segmentally labeled synuclein, minimal media had the same composition as above, but in addition, the remaining 14 unlabeled amino acids (C, D, E, F, H, I, K, M, N, P, Q, R, W, Y) were added at a final concentration of 100 mg/L. For ^13^C_5_-valine, ^13^C_6_-Leucine-segmentally labeled synuclein, the same M9 salts and minerals were added but 4 g/L of natural abundance glucose and 1 g/L of natural abundance NH_4_Cl were used. Also, ^13^C_5_-valine and ^13^C_6_-Leucine were added at a final concentration of 100 mg/L. The remaining 18 natural abundance amino acids (A, C, D, E, F, G, H, I, K, M, N, P, Q, R, S, T, W, Y) were added at a final concentration of 100 mg/L.

Natural-abundance recombinant Cfa_C_-α-syn(C30-140) was expressed in LB media (10 g/L tryptone, 5 g/L yeast extract, 10 g/L NaCl). Cells were grown at 37 °C with shaking until their *A*_600_ reached 0.6–0.8, and protein expression was induced by the addition of IPTG to a final concentration of 1 mM. Cells were collected 4 h later by centrifugation at 4,000 × *g* for 15 min at 4 °C.

Proteins from constructs 1 and 2 were purified using nickel resin. Cell pellets were lysed by resuspension in 20 mL of binding buffer (100 mM NaH_2_PO_4_, 500 mM NaCl, 4 mM TCEP, pH 8) per liter of growth. Protease inhibitor tablets were added. The cell suspension was sonicated 6 times (pulse: 60% amplitude, 30s, 0.5 s on, 0.5 s off). Lysates were centrifuged for 20 min at 35,000 × *g* to remove insoluble cellular debris. Cleared lysates were filtered using 0.2 µM filter then passed through a 15-mL Nickel column using Biorad NGC system. The column was washed with 10-15 column volumes of binding buffer (100 mM NaH_2_PO_4_, 500 mM NaCl, 2 mM TCEP, pH 8). It was then washed with 10-15 column volumes of wash buffer (100 mM NaH_2_PO_4_, 500 mM NaCl, 10 mM imidazole, 2 mM TCEP, pH 8). The protein was eluted in elution buffer (100 mM NaH_2_PO_4_, 500 mM NaCl, 450 mM imidazole, 2 mM TCEP, pH 8). Purity was assessed by SDS/PAGE. Protein concentration was quantitated using theoretical extinction coefficients of 26930 M^−1^ cm^−1^ for construct 1 and 7450 M^−1^ cm^−1^ for construct 2.

### Split-Intein Ligation and Purification

The purified split-intein constructs were ligated with a molar excess of 1:1.5 (construct 1:construct 2) in ligation buffer [50 mM Tris, 300 mM NaCl, 2 mM Tris(2-carboxyethyl)phosphine (TCEP), 0.5 mM EDTA, pH 7.4]. The final concentrations of constructs 1 and 2 were 25 µM and 30 µM, respectively. The reaction mixture was left at 4 °C overnight. It was then concentrated 5-fold using a 10-kDa molecular mass cutoff Amicon spin concentrator. The reaction mixture was then incubated with Ni-NTA resin for an hour at room temperature.

The resin was transferred to a PD-10 column and washed with 1 column volume of binding buffer. The flowthrough and the wash were collected since they contain the untagged α-syn. The protein solution was concentrated to around 1 mM and buffer exchanged into a 6 M guanidine hydrochloride, 50 mM sodium phosphate, pH 7.2 buffer suing PD10 columns before proceeding to the desulfurization step.

Desulfurization was done following a previously published protocol, which contains important safety precautions (https://link.springer.com/protocol/10.1007/978-1-49-2978-8_1#Sec2). Briefly, pH-adjusted TCEP was added to the protein solution to bring up the concentration of TCEP to 100 mM. Then the radical initiator solution (100 mM VA-044 in 6 M guanidine hydrochloride, 50 mM sodium phosphate, pH 7.2) was added to a final concentration of 6 mM. The following steps are all done in a fume hood. Methylpropane-2-thiol (or t-butyl mercaptan) was added in the hood to a final concentration of 400 mM. The reaction was left on an Eppendorf orbital shaker for 2 hours at 600-800 rpm at 37 °C. The protein was then isolated in the hood using PD10 columns equilibrated with the buffer of interest.

At this step we had α-syn and an 11-kDa byproduct of the ligation reaction (the last 111 amino acids of α-syn). Therefore, after the desulfurization reaction, the protein was concentrated and loaded onto a Superdex 75 size exclusion column. The fractions containing the highest concentration of α-syn and the lowest concentration of the 11-kDa species were pooled together and concentrated.

#### DNP sample preparation

The ^13^C, ^15^N-segmentally labeled synuclein and VGLAST-segmentally labeled α-syn monomer in buffer samples were prepared by buffer exchanging the proteins into a 50 mM sodium phosphate, 0.1 mM EDTA, 0.02% (*w/v*) sodium azide pH 7.4 buffer with 88% deuteration. Then each protein was concentrated to around 900 µM. ^12^C_3_, *d_8_*-glycerol was added to a final concentration of 15%. AMUPol was added to a final concentration of 5 mM.

The VGLAST monomer in urea sample was prepared by buffer exchanging the protein into a 50 mM sodium phosphate, 8.7 M urea, pH 7.4 buffer with 55% deuteration. Then it was concentrated to around 900 µM. ^12^C_3_, *d_8_*-glycerol was added to a final concentration of 15%. AMUPol was added to a final concentration of 5 mM. The final deuteration level of this sample is 44% (not 75%) because of the very high concentration of urea.

The 2N0A fibril samples of ^13^C, ^15^N-segmentally labeled α-syn and ^13^C_5_-valine, ^13^C_6_-Leucine-segmentally labeled α-syn were prepared as described (23, 24). Briefly, we buffer exchanged the monomer samples into a 50 mM sodium phosphate, 0.12 mM EDTA, 0.02% sodium azide (*w/v*), pH 7.4 buffer. The monomer samples were then each concentrated to 1 mM and incubated at 37 °C and 200 rpm (orbital shaker, Eppendorf) for 3 weeks. The fibrils were then pelleted using an ultracentrifuge at 200,000 x g for 40 minutes. The pellet (□ 40 mg) was washed by resuspending it in 1 mL of 88% deuterated PBS. The suspension was centrifuged again at 200,000 x g for 40 minutes. After calculating how much ^12^C_3_, *d_8_*-glycerol and AMUPol is needed to achieve final concentrations of 15% glycerol and 7 mM AMUPol, the glycerol/AMUPol mixture was placed on the opposite side of the fibril pellet in the ultracentrifuge tube, then centrifuged at 200,000 x g for 15-20 minutes. This process was repeated two more times with rotating the tube each time in order to incorporate the glycerol/AMUPol mixture into the fibril pellet. The pellet was scooped and packed into 3.2 mm sapphire rotors using a table-top centrifuge. Samples were stored frozen.

#### NMR experiments and procession

All dynamic nuclear polarization magic angle spinning nuclear magnetic resonance (DNP MAS NMR) experiments were performed on a 600 MHz Bruker Ascend DNP NMR spectrometer/7.2 T Cryogen-free gyrotron magnet (Bruker), equipped with a ^1^H, ^13^C, ^15^N triple-resonance, 3.2 mm low-temperature (LT) DNP MAS NMR Bruker probe (600 MHz). The sample temperature was 104 K, and the MAS frequency was 12 kHz. ^13^C–^13^C 2D correlations were measured by using 10 ms DARR mixing. A total of 256 complex *t*_1_ points with an increment of 25 μs were recorded. For 2D experiments, DARR data were processed by using NMRPipe. The data were apodised with a Lorentz-to-Gauss window function with an Inverse Exponential width of 25 Hz and a Gaussian Broaden width of 150 Hz in the *t*_1_ and *t*_2_ time domains.

#### Normalization of spectra

For comparison of the alanine and glycine peaks across the different samples (monomer in buffer vs. monomer in urea vs. 2N0A fibril), the alanine C_α_-C_β_ peak was integrated in each of the following samples: ^13^C, ^15^N-segmentally labeled α-syn, VGLAST-segmentally labeled α-syn and ^13^C,^15^N, segmentally labeled 2N0A fibril. The intensity of that alanine peak across the 3 different spectra was matched by multiplying each spectrum by the factor needed for matching using the nmrPipe command: “nmrPipe -fn MULT -c “constant” -in “sample name”.ft2 -out “adjusted sample name”.ft2 -ov”

#### Generation of predicted peak shapes from statistical coil ensembles

The methods of generating the 10,000 pose poly-alanine host guest peptide ensembles were followed according to Kragelj et al., 2023(37). The predicted peak shapes were made through simulating a random coil of a poly-alanine host-guest peptide. Provided the sequence, we used Flexible Meccano (54) to generate 10,000 PDB structures of a disordered polypeptide chain. In the building process, the phi/psi angles that constitute the backbone of the peptide are sampled from a restricted coil library to provide a statical based sampling approach. Then, the fixbb program from Rosetta Commons optimizes sidechain-rotamer on the fixed backbone previously generated from Flexible Meccano. Finally, we use PPM_One (55) to predict the chemical shifts from the final .pdb file output from Rosetta Commons. To visualize the data, a 2d histogram was made using numpy and 1.5 ppm of line broadening was added. The text file was converted to a .ucsf file format using the “con2Dtext.com” command in NMRpipe to generate a .ucsf file, which was analyzed in Poky (56).

#### Estimation of population and error

Cross-peaks were fit to one to four Gaussian peaks using Poky (56) peak integration settings that allowed for constrained peak motions. Peaks above and below the diagonal were fit to the same number of Gaussians. Results of individual fits are reported in Supplemental Table 1.

Populations of different conformational states were determined independently for the peaks above and below the diagonal and reported as the average ± the standard deviation.

