## Supplementary material for "Structural context modulates the conformational ensemble of the intrinsically disordered amino terminus of α-synuclein": Figure S1, Figure S2, Table S1

**The Supplementary information file includes:**

Figures S1 and S2

Table S1

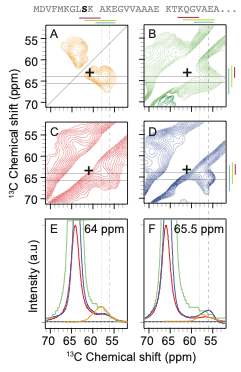

***Figure S1****: Context alters the conformational preferences of serine residues in the amino terminal intrinsically disordered region of α-syn. The primary sequence of the isotopically labeled segment of α-syn is shown with the serine residue highlighted in black. A) The predicted peak shape for the Cα-Cβ cross peak of serine from a statistical coil structural ensemble assuming a homogenous line width of 1.5 ppm. ^13^C-^13^C DARR spectra of frozen segmentally isotopically labeled α-syn in B) 8 M urea C) in the monomeric form in buffer and D) in the amyloid fibril form in buffer. Colored bars annotate the average chemical shift ± two standard deviations for α-helices (magenta), random coils (light green) and β-strands (light blue). The + symbol marks the center of the α-helical cross-peak region. Spectra were normalized to have the same integrated intensity. Horizontal lines at 64 ppm and 65.5 ppm mark the location of 1D slices in E (64 ppm) and F (65.5 ppm). E) Overlay of the normalized one-dimensional slices from the 2D DARR spectra at 64 ppm. F) Overlay of normalized one-dimensional slices from the 2D DARR spectra at 65.5 ppm. Samples contained 15% d_8_-glycerol and 5 mM AMUPol. Spectra were collected at 600 MHz with 12.5 kHz MAS at 104 K.*

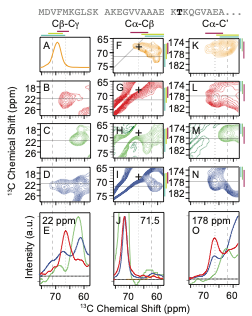

***Figure S2****: Context alters the conformational preferences of T22 in the amino terminal intrinsically disordered region of α-syn. The primary sequence of the isotopically labeled segment of α-syn is shown with the T22 highlighted in black. A) The predicted peak shape for the Cβ of threonine from a statistical coil structural ensemble assuming a homogenous line width of 1.5 ppm. The Cβ-Cg cross-peaks from the ^13^C-^13^C DARR spectra of frozen segmentally isotopically labeled α-syn in B) in the monomeric form in buffer C) 8 M urea and D) in the amyloid fibril form in buffer. Colored bars annotate the average chemical shift ± two standard deviations for α-helices (magenta), random coils (light green) and β-strands (light blue). Spectra were normalized to have the same integrated intensity. Horizontal line marks the location of 1D slices in E. E) Overlay of the one-dimensional slices from the 2D DARR spectra at 22 ppm. F) The predicted peak shape for the Cα-Cβ cross-peak of threonine from a statistical coil structural ensemble assuming a homogenous line width of 1.5 ppm. ^13^C-^13^C DARR spectra of frozen segmentally isotopically labeled α-syn in G) in the monomeric form in buffer H) 8 M urea and I) in the amyloid fibril form in buffer. Horizontal line marks the location of 1D slices in J. J) Overlay one-dimensional slices from the 2D DARR spectra at 71.5 ppm. K) The predicted peak shape for the Cα-CO cross-peak of threonine from a statistical coil structural ensemble assuming a homogenous line width of 1.5 ppm. ^13^C-^13^C DARR spectra of frozen segmentally isotopically labeled α-syn in L) in the monomeric form in buffer M) 8 M urea and N) in the amyloid fibril form in buffer. Horizontal lines at 178 ppm mark the location of 1D slices in J. O) Overlay of the normalized one-dimensional slices from the 2D DARR spectra at 178 ppm. Samples contained 15% d_8_-glycerol and 5 mM AMUPol. The + symbol marks the center of the α-helical cross-peak region. Spectra were collected at 600 MHz with 12.5 kHz MAS at 104 K.*

| **ALANINE** |  | **peak center (ppm)** | |  |  | **peak width (ppm)** | | **population** |
| --- | --- | --- | --- | --- | --- | --- | --- | --- |
| **Condition** | **diagonal** | **C**α | **C**β | **Volume** | **SNR** | **C**α | **C**β | **(%)** |
| predicted | n/a | 55.3 | 18.0 | 16 | n/a | 2.1 | 1.4 | 16 |
| predicted | n/a | 53.2 | 18.6 | 60 | n/a | 2.7 | 2.1 | 60 |
| predicted | n/a | 51.3 | 21.0 | 15 | n/a | 2.6 | 3.2 | 15 |
| predicted | n/a | 53.3 | 16.0 | 8 | n/a | 3.3 | 1.9 | 8 |
| 8 M urea | above | 52.4 | 18.5 | 296 | 17 | 4.0 | 3.4 | 100 |
| 8 M urea | below | 52.4 | 18.6 | 424 | 25 | 4.0 | 3.0 | 100 |
| monomer | above | 55.0 | 18.4 | 265 | 138 | 4.0 | 2.7 | 68 |
| monomer | below | 55.0 | 18.5 | 142 | 74 | 4.0 | 2.9 | 67 |
| monomer | above | 52.2 | 19.4 | 127 | 56 | 4.0 | 3.2 | 32 |
| monomer | below | 52.1 | 19.3 | 70 | 33 | 4.0 | 3.2 | 33 |
| fibril | above | 55.1 | 18.2 | 252 | 33 | 4.0 | 1.7 | 28 |
| fibril | below | 55.0 | 18.8 | 330 | 44 | 3.1 | 2.0 | 29 |
| fibril | above | 52.6 | 18.5 | 266 | 19 | 3.7 | 3.0 | 30 |
| fibril | below | 52.0 | 19.3 | 442 | 44 | 4.0 | 3.2 | 38 |
| fibril | above | 50.8 | 23.1 | 378 | 26 | 4.0 | 2.8 | 42 |
| fibril | below | 50.7 | 23.5 | 378 | 35 | 4.0 | 2.3 | 33 |

| **GLYCINE** |  | **peak center (ppm)** | |  |  | **peak width (ppm)** | | **population** |
| --- | --- | --- | --- | --- | --- | --- | --- | --- |
| **Condition** | **diagonal** | **C**α | **CO** | **Volume** | **SNR** | **C**α | **CO** | **(%)** |
| predicted | n/a |  |  | 100 | n/a | 2.7 | 3.1 | 100 |
| 8 M urea | above | 44.5 | 173.9 | 532 | 24 | 4.0 | 4.0 | 100 |
| 8 M urea | below | 44.6 | 174.6 | 217 | 11 | 4.0 | 4.0 | 100 |
| monomer | above | 47.0 | 177.6 | 262 | 154 | 3.3 | 3.8 | 46 |
| monomer | below | 47.0 | 177.6 | 142 | 76 | 3.7 | 3.4 | 48 |
| monomer | above | 45.4 | 174.6 | 267 | 90 | 4.0 | 4.0 | 47 |
| monomer | below | 45.7 | 174.8 | 123 | 45 | 3.9 | 4.0 | 42 |
| monomer | above | 43.6 | 171.7 | 40 | 90 | 3.2 | 2.7 | 7 |
| monomer | below | 44.0 | 172.1 | 30 | 39 | 3.4 | 3.0 | 10 |
| fibril | above | 46.7 | 177.7 | 333 | 38 | 2.8 | 2.3 | 24 |
| fibril | below | 46.9 | 177.4 | 178 | 15 | 4.0 | 2.4 | 29 |
| fibril | above | 44.9 | 173.8 | 768 | 41 | 4.0 | 3.5 | 56 |
| fibril | below | 45.2 | 173.6 | 338 | 17 | 4.0 | 4.0 | 54 |
| fibril | above | 44.9 | 170.6 | 274 | 46 | 2.1 | 2.2 | 20 |
| fibril | below | 44.9 | 170.6 | 105 | 23 | 2.1 | 2.2 | 17 |

| **LEUCINE** |  | **peak center (ppm)** | |  |  | **peak width (ppm)** | | **population** |
| --- | --- | --- | --- | --- | --- | --- | --- | --- |
| **Condition** | **diagonal** | **C**α | **C**β | **Volume** | **SNR** | **C**α | **C**β | **(%)** |
| predicted | n/a | 56.9 | 41.2 | 8 | n/a | 2.3 | 1.5 | 8 |
| predicted | n/a | 54.5 | 41.6 | 64 | n/a | 3.0 | 2.2 | 64 |
| predicted | n/a | 52.4 | 42.7 | 21 | n/a | 2.3 | 1.5 | 21 |
| predicted | n/a | 53.7 | 39.2 | 7 | n/a | 2.7 | 1.9 | 7 |
| 8 M urea | above | 54.8 | 41.7 | 66 | 4 | 4.0 | 2.6 | 100 |
| 8 M urea | below | 54.6 | 42.2 | 74 | 4 | 4.0 | 3.6 | 100 |
| monomer | above | 57.4 | 41.0 | 49 | 20 | 3.7 | 4.0 | 41 |
| monomer | below | 57.3 | 41.6 | 47 | 20 | 3.5 | 4.0 | 39 |
| monomer | above | 54.2 | 42.6 | 72 | 23 | 4.0 | 4.0 | 59 |
| monomer | below | 53.8 | 42.7 | 72 | 23 | 4.0 | 4.0 | 61 |
| fibril | above | 57.4 | 42.4 | 44 | 44 | 2.8 | 4.0 | 28 |
| fibril | below | 57.5 | 41.3 | 51 | 47 | 2.6 | 4.0 | 29 |
| fibril | above | 54.1 | 42.6 | 47 | 37 | 3.7 | 4.0 | 31 |
| fibril | below | 54.2 | 42.6 | 54 |  |  |  | 31 |
| fibril | above | 53.7 | 46.4 | 64 | 54 | 4.0 | 3.4 | 41 |
| fibril | below | 53.7 | 46.5 | 72 | 69 | 4.0 | 2.6 | 40 |

***Table S1****: Chemical shifts of the center of the fit Gaussian line shape, peak volumes, peak signal to noise ratios, peak widths, and relative peak populations for alanine, glycine and leucine cross peaks in the predicted peak for a statistical coil ensemble, for the experimental peaks both above and below the diagonal in the experimental spectra for the amino terminal region of a-syn in 8 M urea, as a monomer in buffer and in the disordered region that flanks the amyloid core.*
